# Atypical septate junctions maintain the somatic enclosure around maturing spermatids and prevent premature sperm release in *Drosophila* testis

**DOI:** 10.1101/360115

**Authors:** Pankaj Dubey, Tushna Kapoor, Samir Gupta, Seema Shirolikar, Krishanu Ray

## Abstract

Tight junctions prevent the paracellular flow and maintain cell polarity in an epithelium. These are also essential for maintaining the blood-testis-barrier involved in regulating sperm differentiation. Septate junctions are orthologous to the tight junctions in insects. In *Drosophila* testis, major septate junction components co-localize at the interface of germline and somatic cells initially and then condense between the two somatic cells in a cyst after germline meiosis. Their localization is extensively remodeled in subsequent stages. We find that characteristic septate junctions are formed between the somatic cyst cells at the elongated spermatid stage. Consistent with the previous reports, knockdown of essential junctional components, Discs-large-1 and Neurexin-IV, in the somatic cyst cells, during the early stages, disrupted sperm differentiation beyond the spermatocyte stage. Somatic knockdown of these proteins during the final stages of spermatid maturation caused premature release of spermatids inside the testes, resulting in partial loss of male fertility. These results indicate the importance of maintaining mechanical integrity of the somatic enclosure during spermatid coiling and release in *Drosophila* testis. It also highlights the functional similarity with the tight junction proteins during spermatogenesis in mammalian testes.

**Summary statement:** Dubey et al., showed that septate junctions stitch the somatic enclosure around maturing spermatids in *Drosophila* testis. Maintaining the integrity of this junction is essential for proper release of spermatids.

## Introduction

Germ cell development requires appropriate microenvironment. In male germline, it is provided by the somatic-origin cells, *viz.*, the Sertoli cells in mammals, and the somatic cyst cells (SCCs) in *Drosophila* (Griswold, 1998; Zoller and Schulz, 2012). Both these cell types insulate developing germ cells from the body fluids, and thus, from the immune system. In mammals, this is accomplished by a specialized structure called the blood-testis-barrier (BTB) (Cheng and Mruk, 2012). Tight junctions (TJ) form an essential part of the BTB (Mruk and Cheng, 2015). In an epithelium, TJs restrict the paracellular flow of solutes from the lumen, as well as separate the apical and basolateral domains of the plasma membrane (Hartsock and Nelson, 2008). In testis, TJs between Sertoli cells at the BTB play a significant role in maintaining seminiferous tubule architecture, as well as the progression of spermatogenesis. In mice testis, knockout of Claudin-11 (Cldn11), an essential component of tight junctions in the testis, led to detachment of Sertoli cells from the basement membrane, thereby severely affecting the progression of spermatogenesis and the reproductive output (Mazaud-Guittot et al., 2010). Loss of another tight junction protein, Zona-occludens-2, from the Sertoli cells leads to mislocalization of a number of BTB proteins such as Cldn11, the gap junction protein-Connexin-43, and actin, which leads to loss of BTB integrity, and a decrease in male fertility (Xu et al., 2009). Together, these observations suggested that TJs maintain the integrity of the seminiferous tubule and the BTB.

Septate Junctions (SJs) in insects are considered to be the functional equivalent and evolutionary precursor to TJs (Banerjee et al., 2006). Both of these junctions require the Claudin family of proteins for the formation and maintenance of barrier function (Furuse et al., 1998a; Furuse et al., 1998b; Nelson et al., 2010; Wu et al., 2004). In *Drosophila*, SJs are identified by the typical, ladder-like arrangement of electron-dense elements at ~15 nm interval along the membrane interface between two epithelial cells (Banerjee et al., 2006; Locke, 1965). Structurally, SJs are classified into two types - pleated and smooth. Pleated SJs (pSJs) are found in ectoderm-derived epithelia. The pSJs have a typical ladder-like arrangement of electron-dense septa connecting the plasma membrane pair, while smooth SJs have a parallel arrangement (Banerjee et al., 2006). Neuroglian (Nrg), Neurexin (NrxIV), Na^+^/K^+^-ATPase-α, Nervana (Nrv2), Lachesin (Lac), Disc large 1 (Dlg1), Coracle (Cora), and Fasciclin III (Fas III) constitute SJs in *Drosophila* (Banerjee et al., 2006; Woods et al., 1996). SJs play a critical role in developing tissue architecture and function. The homozygous *nrx* mutants fail to form a blood-nerve barrier due to the disruption of the SJs (Baumgartner et al., 1996). Loss of Dlg1 results in abnormal growth and fusion of imaginal discs (Woods and Bryant, 1989); while the loss of *cora* causes dorsal closure defects (Fehon et al., 1994). Also, N^+^/K^+^ ATPase-α and nervana-2 are involved in tracheal tube size control, independent of their role in maintaining the diffusion barrier (Paul et al., 2003). These results further indicated that SJs are involved in both cell signaling and the maintenance of epithelial integrity.

In *Drosophila* testis, the spermatogonia develop inside an enclosure formed by two somatic-origin cyst cells (SCCs) that undergo extensive morphogenesis and ultimately differentiate into the Head (HCC) and Tail (TCC) cyst cells during spermatid elongation (Lindsley & Tokuyasu, 1980; White-Cooper, 2004). Each spermatid elongates to ~1.8 mm after meiosis inside the somatic enclosure in the testis. Subsequently, they are individualized, coiled and released into the seminal vesicle (SV) as mature sperm (Lindsley & Tokuyasu, 1980). As suggested previously (Fairchild et al., 2015), we found that critical components of SJs localize at the interface of SCCs during spermatid elongation. We also found that the junction migrates towards the caudal end of the enclosed spermatid head bundle after individualization, which confirmed an earlier prediction (Tokuyasu et al., 1972). Further, Transmission Electron Microscopy (TEM) analysis suggested that the SJs form between the SCCs after the spermatid individualization. The HCC-TCC association is likely to be subjected to high level of tension during the spermatid elongation, and subsequent differentiation. During this process, SJs could presumably impart mechanical stability balancing the tension at the HCC-TCC interface. We found that loss of Dlg1 in SCCs during the spermatid coiling and maturation could disrupt the localization of SJs components at the HCC-TCC interface and result in premature release of spermatids. Time-lapse imaging further indicated that the spermatids are likely to be released during the cyst rotation in the Terminal Epithelium (TE) region. Altogether, these observations suggest that the SJs between HCC and TCC form after the sperm individualization and that the junction is required to maintain the mechanical integrity of the somatic cyst enclosure during its migration through TE before sperm release.

## Results

To probe for the localization of SJ proteins at the cellular interfaces during spermatogenesis, we carried out a limited screen using protein trap lines and antibody staining of adult testis. We identified the SJs proteins –Nrg, Nrx-IV, Na^+^/K^+^-ATPase-α, Nrv2, and Lac - at the soma-germline interface by using protein trap lines (green, Fig 1). Anti-Dlg-1 staining (red, Fig 1) of these testes preparations suggested that all the above SJ components colocalize with Dlg1 at all stages. Further, the pattern matched with that of the endogenous Dlg1-GFP (red, Fig 1) and Cora immunostaining (green, Fig 1). These SJ components localized at the germ-soma interface during the early stages, until the completion of meiosis (Fig 1A). We also found a condensed and prominent localization near the caudal end of the spermatid head bundles during the coiled stages, in the TE region (Fig 1B). Together, these results indicated that the cellular interface marked by these proteins undergoes extensive reorganization between the early and late stages of spermatogenesis.

**Fig 1.**
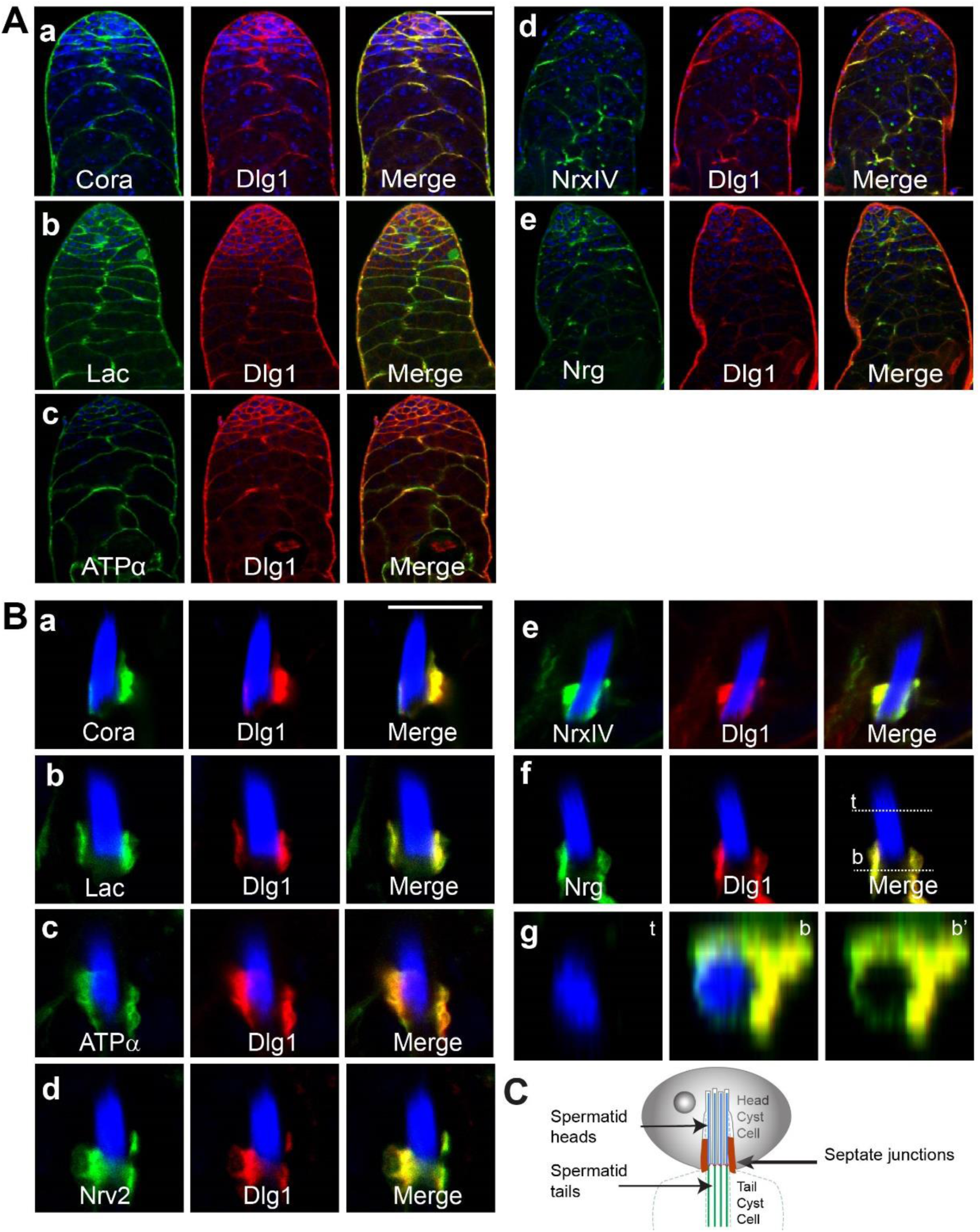
Major components of the septate junction co-localize during early and late stages of spermatogenesis. A. Apical tips of testes showing co-localization of the SJ proteins - Lac-GFP **(b)**, ATPα-GFP **(c)**, Nrx-IV-GFP **(d)** and Nrg-GFP **(e)** - with Dlg1 (red) immunostaining at the interface of the germline and somatic cells. Testis from the Dlg1GFP (red) stock was immunostained with anti-Cora (green) **(a)**. All specimen were stained with the Hoechst dye (blue) marking the nuclei. (Scale-50 μm)
B. The SJ proteins also localize near the compact nuclei bundle (NB) of the mature spermatids during the late stages. Hoechst staining, marking all nuclei, is in blue. **(g)** depicts X-Z digital section through the top (t) and bottom (b) parts of the specimen shown in f, indicating that the SJ proteins localize all around the NB. (Scale-10 μm)
C. Schematic describes the position of the junction, between the head and tail cyst cells.

### Morphogenesis of the cellular interfaces marked by SJs proteins during spermatogenesis

Next, we followed the morphogenesis of the SCC interfaces, using Nrg-GFP protein-trap, throughout all the stages of spermatogenesis (Fig 2A). Consistent with a previous report (Papagiannouli and Mechler, 2009), which analyzed the localization of Dlg, Nrg-GFP was found around all germ cells and SCCs at the early spermatogonial stages (Fig 2B, arrowheads and arrows). Nrg-GFP was mostly enriched around the entire cyst during the late spermatogonial (arrows, Fig 2B) and post-meiotic spermatocyte stages (arrows, Fig 2C), indicating a gross reorganization of the junctions during these stages. The elongating stage cyst can be identified by polarized localization of sixty-four spermatid nuclei on one side and Spectrin caps (identified by α–spectrin immunostaining) on the other (Ghosh-Roy et al., 2004). Squash preparation of testis revealed localization of Nrg-GFP at the middle of the early elongating (Fig 2D), as well as in fully elongated cysts (Fig 2E). Consistent with an earlier report (Fairchild et al., 2015), these observations suggest that SJ proteins localize at the HCC-TCC boundary from the elongation stages onwards.

**Fig 2.**
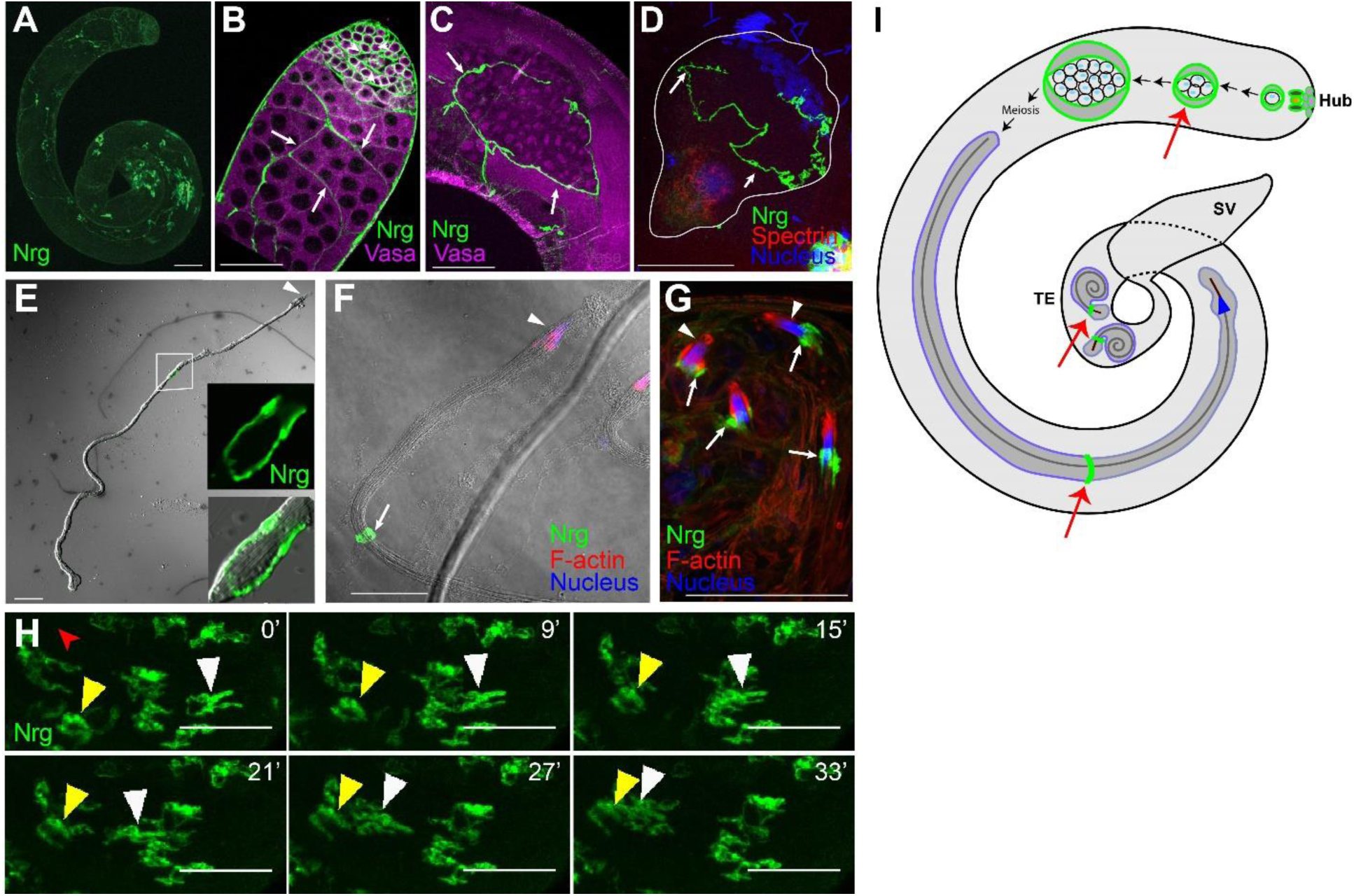
Morphogenesis of the SJ protein Nrg during spermatogenesis. **A-C)** Two-day-old Nrg-GFP (green) testes stained with the anti-Vasa antibody (magenta). **(A)** Low magnification image of Nrg-GFP testis shows the presence of Nrg at different stages. **(B)** Apical end of the testis shows Nrg-GFP localization around individual spermatogonia (arrowheads) at the initial stage. It is then restricted to the cyst perimeter (arrows) of the primary spermatocyte stages. **(C)** A post-meiotic cyst shows the presence of Nrg-GFP along the cyst perimeter. Nrg-GFP is excluded from the germ cell perimeter inside the cyst enclosure from the spermatocyte stage onwards. **D-F)** Squash preparation Nrg-GFP testes immunostained with the anti-Spectrin antibody (red, **D**), Hoechst dye (blue) and Phalloidin (red; **F**). **(D)** An early elongating cyst (outlined by white boundary) shows polarization of the spermatid nuclei (blue) and tails (red), and localization of Nrg-GFP (arrows) at the HCC-TCC interface. **(E-F)** Elongated spermatid cysts from Nrg-GFP testis were isolated and stained for the F-actin cone (red) and nucleus (blue) marking the rostral ends (arrowheads). The HCC and TCC are highly extended at this stage, and a condensed form of Nrg-GFP (arrows) between these two cells was seen in the middle region. **G)** Coiled stage spermatids from Nrg-GFP testis, stained for F-actin (red) and nucleus (blue). The arrow indicates localization of the Nrg (arrow) near the spermatid nuclei bundle (arrowhead, blue). Note that the position of the Nrg-GFP has changed post individualization. **H)** Time-lapse images of Nrg-GFP testis shows the movement of Nrg-GFP structure (yellow and white arrowheads) towards basal end of the testis. The red arrowhead shows the location of the SV. (Scale-50 μM for all panels) **I)** Schematic illustrates the morphogenesis of domains marked by SJ proteins in adult testis. Schematic is not to scale.

The compact localization of Nrg-GFP around the middle of the cyst persisted until the beginning of the individualization stage (arrowhead, Fig 2F). Post individualization, Nrg-GFP was found at the caudal end of compacted spermatid nuclei bundle (NB) (arrows, Fig 2G). Thus, the junction appeared to move towards the NB during individualization or early coiling stage. Time-lapse imaging in the mid-region of testis further indicated that occasionally an SJ moves towards the base (yellow and white arrowheads, Fig 2H; Movie S1). Nuclei bundles (NBs) of elongated spermatids within a cyst are positioned near the base of the testis and a compacted SJ forms in the middle of these cysts (Figure 2I). The spermatid tails coil up after individualization, and the SJ is repositioned near the rostral end of the NB between the HCC and TCC. Therefore, the SJ movement towards the base may suggest a reorganization of the HCC and TCC morphology, either before or during coiling. Previously, TEM studies predicted that the junction is repositioned during individualization or late coiling stage and that this movement coincides with the condensation of the HCC towards the spermatid head bundle and expansion of TCC to cover the entire tail bundle (Tokuyasu et al., 1972). Our results provide experimental proof of this previously proposed model.

### SJs were first observed between the somatic cyst cells during the elongated spermatid stage

We further examined the cellular interfaces using TEM in wild-type testis. The germline and somatic cell interfaces during the pre-elongation stages revealed no specific electron-dense structures (Fig 3A-A’). We were not able to identify any SJs during pre-meiotic stages. Some electron-dense structures were seen between the SCCs around the elongated spermatids (arrow, Fig 3B-B’). More prominent electron dense patterns, resembling the septate junctions, were found around the tails of fully elongated spermatids containing major and minor mitochondrial derivatives around the axoneme (arrows, Fig 3C-C’). These electron-dense SJs were also found near the nuclei of compacted spermatid head bundles at the base of the testis, which is characteristic of the post-individualized stages (arrow, Fig 3D-D’). Together with the previous results (Tokuyasu et al., 1972), these observations further suggested that SJs are formed between the somatic cells, the HCC and TCC, during spermatid elongation and maintained in subsequent stages.

**Fig 3.**
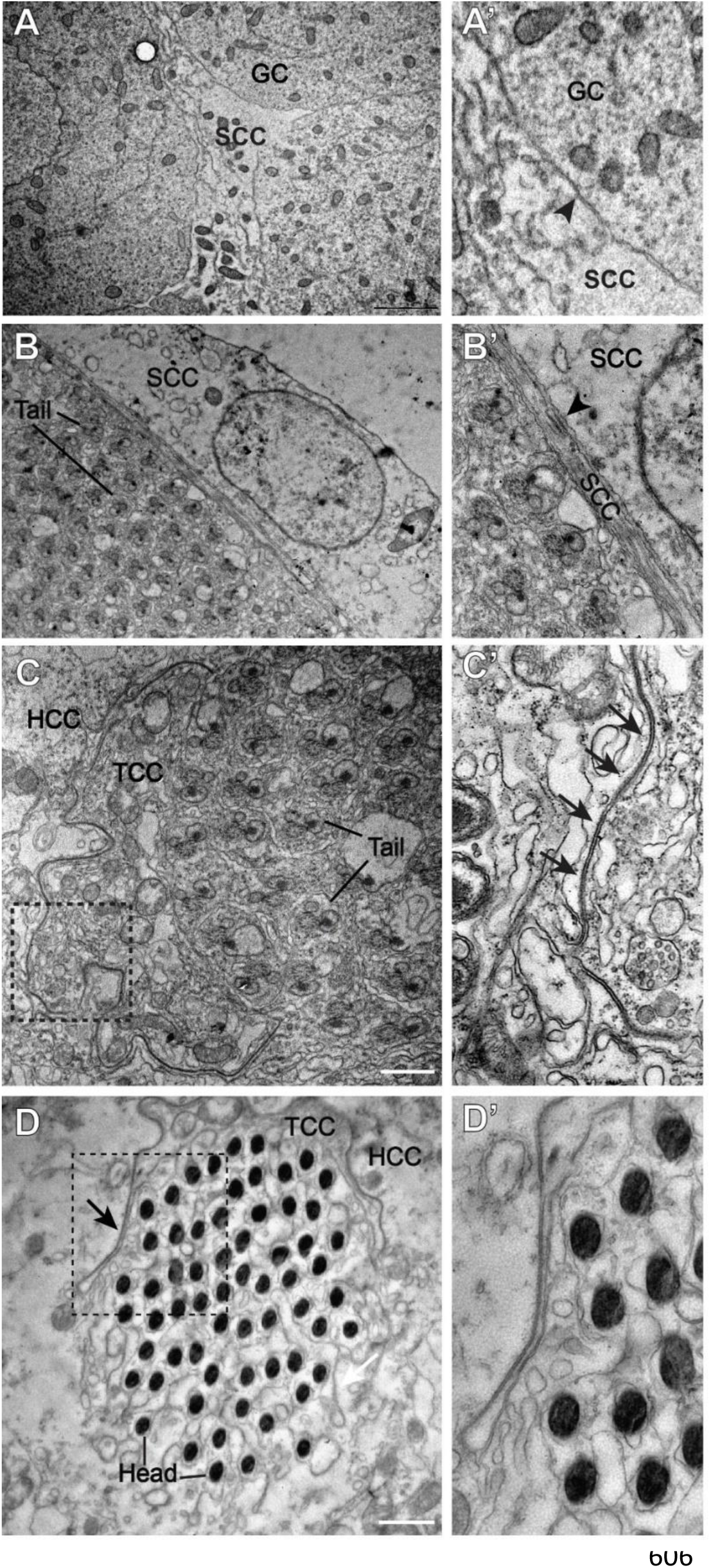
Ultrastructural analysis of the germ-soma and soma-soma interfaces in adult testis. A. Electron micrograph of the spermatogonia! stages shows a somatic cyst cell (SSC) and germ cells (GC). The junction between the SSC and GC (arrowhead) is seen (**A’**).
B. Section through an elongated cyst, as can be identified by the spermatid tails with major and minor mitochondria, along with associated SSC. Note that the junction between the SSCs does not resemble an SJ (arrowhead, **B’**).
C. Electron micrograph through the tails of a more mature, pre-individualized cyst shows the presence of a septa-like pattern between the two surrounding cyst cells. **C’** shows the magnified image of the boxed region in **C**. Arrows indicate a clear ladder-like septate junction between the membranes of the two somatic cyst cells.
D. Section through the spermatid heads at the coiled stages, with surrounding HCC and TCC. **D’** shows a magnified image of the boxed region in **D**. Ladderlike arrangement of SJ between the two cyst cells can be seen around the sperm head. Note that similar to the results obtained by confocal microscopy, the junction has relocated just caudal to the sperm heads. (Scale-1 μM)

### Knock-down of Dlg1 and Nrx-IV in the somatic cyst cells at an early stage arrested post-meiotic differentiation

In adult *Drosophila* testis, Cora and Nrx IV in the SCCs are essential for forming a functional germ-soma permeability barrier and for further germline differentiation (Fairchild et al., 2015). However, as discussed earlier, SJ proteins also have roles other than serving as a diffusion barrier. For instance, a significant number of pole cells in the *Dlg1* homozygous mutant embryos fail to reach the gonadal pockets. Further, the male-specific mesoderm cells expressing Sox100B fail to get incorporated into the male gonad in stage-15 embryos (Papagiannouli, 2013). In *Dlg1* homozygous mutant larvae, the Eyes-absent (Eya)-positive SCCs were reduced, and apoptosis was induced in the 16-cell spermatocyte cysts, indicating a role of Dlg in somatic differentiation, as well as for the survival and differentiation of germ cells (Papagiannouli and Mechler, 2009). Although the *tj*-*Gal4* mediated knockdown of Dlg1 in the SCCs during the spermatogonial stages disrupted the cyst permeability barrier and arrested differentiation, it did not affect the transit amplifying divisions of the spermatogonia (Gupta et al., 2018). Hence, we conjectured that in addition to maintaining the barrier function, the somatic Dlg1 activity might specifically regulate the transition to the meiotic stages in the male germline.

To further understand the role of the SJs during the spermatogonia to spermatocyte transition, we knocked down two essential components of the junction, Dlg1 and Nrx-IV using *eya*-*Gal4*, which expresses in both the somatic cyst cells from the 4-cell spermatogonial stage onwards (Fig S1A, A’) (Fabrizio et al., 2003; Leatherman and Di Nardo, 2008). The *eya*-*Gal4*> *dsGFP* testis contained tightly packed, mitotically-active, spermatogonial cells with condensed chromatin at the apex (arrows, Fig 4A, B). The chromatin was decondensed at the subsequent spermatocyte stages (arrowheads, Fig 4B). In the *eya*-*Gal4*> *dsDlg1* testes, the apical ends of testes appeared shrunk (Fig 4D-E), and Dlg1 immunostaining was limited to the germline cells (Fig 4E’). The testis was mostly filled with germ cells having compact chromatin morphology (Fig 4E). It was difficult to distinguish individual cysts in these testes, and there were very few elongated spermatids, as compared to control (Fig 4C, F). The Eya immunostaining, however, appeared in the SCCs at appropriate region of the testis (Fig 4C, F’), indicating that loss of Dlg1 may not affect the *eya* expression. In contrast, immunostaining with the other somatic marker Traffic-jam (TJ), which is expressed in the early population of somatic cyst cells in the control testis (Fig 4G-H) (Hudson et al., 2013; Li et al., 2003), revealed abnormal expansion of the staining in the Nrx-IV knockdown testes (Fig 4I-J). Together, these results confirmed that loss of Dlg1 and Nrx-IV from the SCCs during the mitotic-meiotic transition also blocks differentiation beyond the early spermatogonial stages.

**Fig 4.**
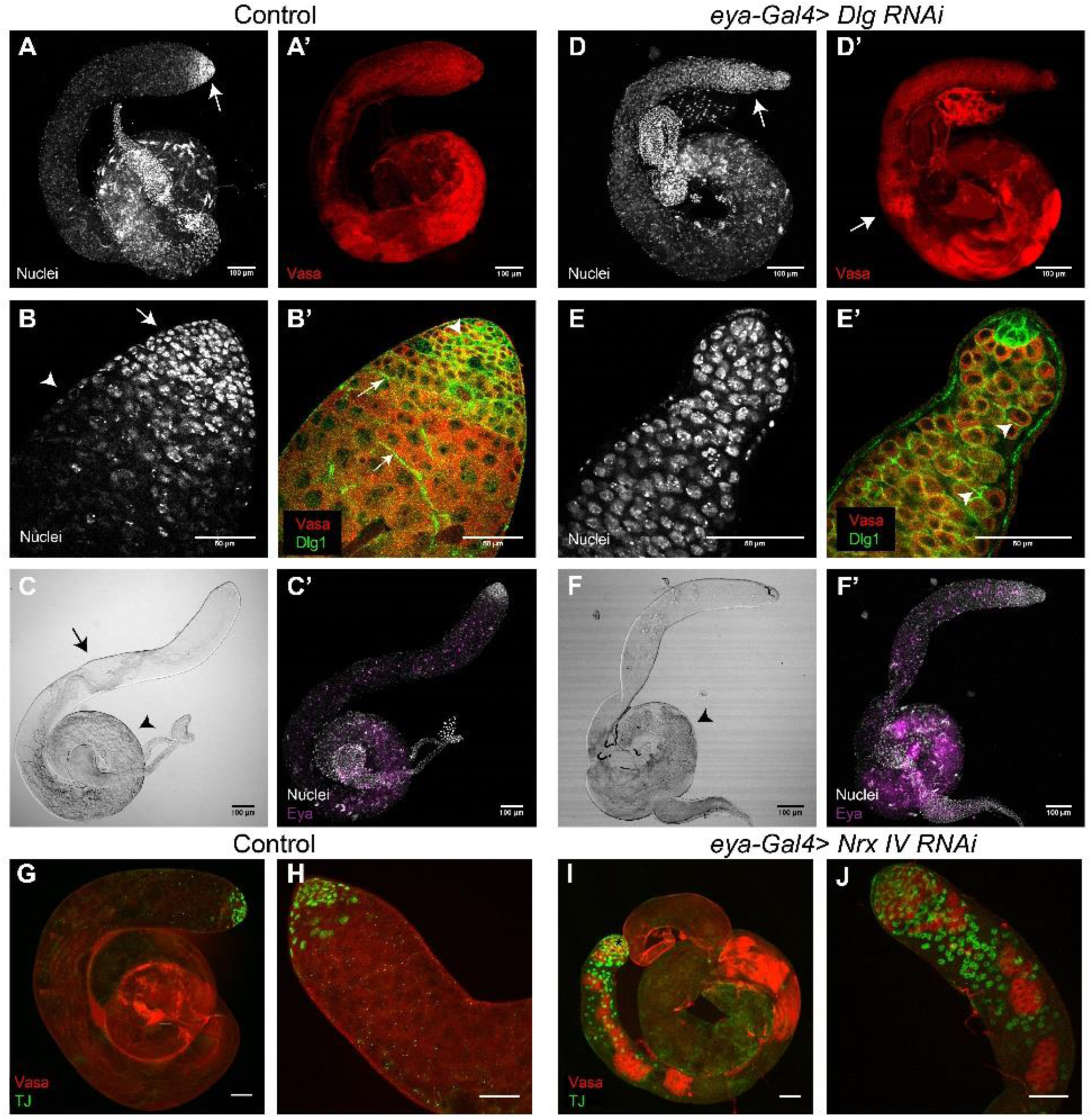
Knockdown of Dlg 1 and Nrx-IV during spermatogonial stages leads to a defect in proliferation and differentiation. **A-B’)** Control (*eya*-*Gal4*> *dsGFP*) testis stained with the Hoechst dye (white), Dlg1 (green) and Vasa (red). **(A)** Hoechst staining shows tightly packed, condensed nuclei at the apical tip (arrow). **(A’)** Vasa pattern in control testis. **(B)** High magnification image of the apical tip shown in **(A)**. Arrow marks the condensed nuclear staining of mitotically active cells, while arrowhead marks the transition to meiotic stages, as indicated by comparatively less intense nuclear staining. **(B’)** Dlg 1 (green) localizes at the membranes of the SSCs, surrounding the germ cells, as indicated by Vasa (red). **C)** DIC image indicates the presence of elongated/ individualizes tails (arrow), as well as coiled tails (arrowhead). **C’)** Control testis shows the distribution of eya (magenta) positive somatic cyst cells. **D-E’)** *eya*-*Gal4*> *Dlg RNAi* testis stained with the Hoechst dye (white), Dlg1 (green) and Vasa (red). **(D)** The spatially distinct localization of the mitotic clusters is lost in *eya*-*Gal4*> *dsDlg1* testis. Brightly stained mitotic nuclei can be seen extending until the middle region of the testis. **(D’)** Pockets of Vasa staining, usually restricted more apically, extend until the middle region of the testes (arrow). **(E)** High magnification image of the apical tip of the testis shown in **(D)**. Note that the apical tip looks shrunk, as compared to control testes in **(B)**. **(E’)** Dlg1 staining (green) is lost from around the germ cells (red; Vasa). Arrowhead indicates Dlg staining around the germ cells. **F)** DIC image indicates a lack of elongated/individualized tails and a decrease in the density of coiled tails (arrowhead) **F’)** *eya*-*Gal4*> *Dlg RNAi* testis shows the distribution of eya (magenta) positive somatic cyst cells. Distribution of the eya-positive cells is similar to control. **G-H)** Control (Wild-type) testis stained with anti-Vasa (red) and anti-TJ (green) antibodies. **(G)** The restricted pattern of TJ-expressing somatic cells at the apical tip of the testis (marked by an asterisk). **(H)** High magnification of the apical tip of a control testis. **I-J)** *eya*-*Gal4*> *dsNrx*-*IV* testis stained with anti-Vasa (red) and anti-TJ (green) antibodies. **(I)** Patchy expression of Vasa indicates defects in proliferation. TJ-positive cells are no longer restricted at the apical tip (asterisk) of the testis. **(J)** Higher magnification of the apical tip of *eya*-*Gal4*> *dsNrx*-*IV* testis. Also note that the apical tip appeared shrunk, similar to what was seen upon knockdown of Dlg 1. The scale bars indicate 50 μM unless specified otherwise on the image.

### Knockdown of SJs during the coiling stages disrupted spermatid bundles

To determine the role of the SJs in the post-meiotic stages, we used *PpY*-*Gal4*, which expresses in the somatic cyst cells after meiosis (arrowhead, Fig S1B-B’; Jung et al., 2007;). We found that expression of *PpY*-*Gal4*> *dsDlg1* abolished Dlg1 staining from the HCC-TCC interface in the cysts only in the TE region (Fig S2), suggesting Dlg1 is effectively knocked-down at the terminal stages by the dsRNA expression. Knockdown of Dlg1 also led to the loss of Coracle staining around the compact NBs in the TE region (Fig S2F). These results suggest that SJs may be disrupted during the terminal stage of spermiation due to the *PpY*-*Gal4*-mediated expression of the *dsDlg1.* We found that the number of intact NBs in the TE region were significantly reduced (Fig 5A-C, I) and an unusually large number of free spermatid heads (insets, Fig 5A-C) were found at the base of the testes. Corresponding bright field images revealed improperly coiled spermatid tails (Fig 5D-F). In comparison, the morphology of the early and progressed individualization complexes (ICs) were normal in these testes (Fig S3). The occurrences of early ICs (Fig 5G), as well as the number of mature spermatid head bundles (NBs) outside the TE zone (Fig 5H), considered as the indicators of successful completion of spermatid elongation (Ghosh-Roy et al. 2005), was unaffected. These results indicated that the SJs proteins are required in the SCCs to maintain the spermatids in tightly bundled and coiled form. A similar disruption of the spermatid bundle due to loss of F-actin assembly was shown to affect spermatid release earlier (Dubey et al., 2016).

**Fig 5.**
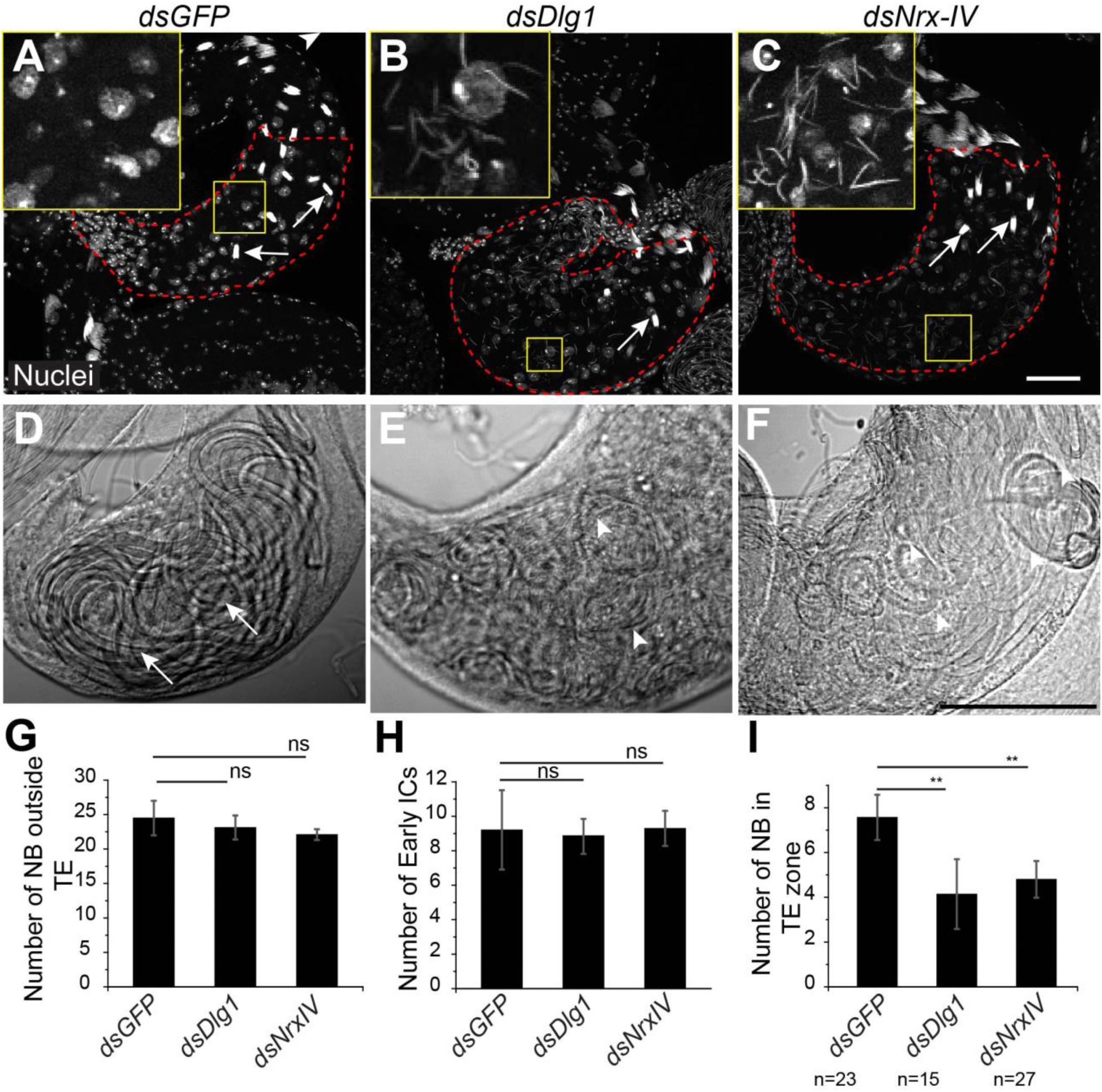
Knockdown of SJ components in the cyst cells during late stages affects NB integrity and spermatid coiling. *A)* Control (*PpY*-*Gal4*> *dsGFP*) testes stained with Hoechst (white) to mark the nuclei. Compact bundles of spermatid nuclei (arrows) are seen in the TE region (red dashed outline). The inset shows that no single spermatid heads can be seen. **B-C)** *PpY*-*Gal4*> *dsDlg1 and PpY*-*Gal4*> *dsNrx*-*IV* testes, respectively, stained with the Hoechst dye. A few intact bundles of spermatid heads (arrows) can be seen in the TE zone (red dashed outline). Unusually large number of disrupted single spermatid heads were found inside these testes (insets). **D-F)** Bright-field images show the basal end of Control **(D)**, *PpY*-*Gal4*> *dsDlg1* (E) and *PpY*-*Gal4*> *dsNrxIV* **(F)** testes. The arrows indicates intact coiled bundle present in the control testis while arrowheads point towards the disrupted bundle in the RNAi backgrounds. **G-I)** Histograms show quantifications of intact NBs (mean±s.d.) outside TE **(G)**, number (mean±s.d.) of early ICs (mean±s.d.) **(H)**, and the number (mean±s.d.) of NBs inside the TE **(I)** in the Control, *PpY*-*Gal4* > *dsDlg1* and *PpY*-*Gal4*> *dsNrxIV* testes. P-value (^⋆⋆^<0.01) was calculated using the Mann-Whitney U test. (Scale-50 μM)

### Knockdown of Dlg1 in SCCs during spermatid coiling induced the premature release of spermatids within the testis

The cyst rotates after entering the TE and the rostral end of the NB of coiled spermatids orients away from the seminal vesicle (SV) at the time of release (Dubey et al., 2016). Time-lapsed imaging also showed that the spermatids are pulled back from the HCC with their tails leading during the release (Movie S2), and the SJs between HCC and TCC remains intact during the release (Dubey et al., 2016). To identify whether the loss of SJs in the SCCs during spermatid coiling could lead to the abnormal release, we estimated the orientation of intact spermatid head bundles (NBs) in the TE region in the *Dlg1* RNAi background. In control testes, NBs found at the 100-200 μm distance from the SV were oriented with equal propensity both towards (arrowheads, Fig 6A) and away (stars, Fig 6A) from the SV (black and grey bars, Fig 6C). In comparison, the majority of the NBs in the 200-300 μm zone were found oriented towards the SV (Fig 6C). In the *Dlg1* RNAi background, a significant fraction of the relatively fewer NBs found in the 100-200 μm zone remained oriented towards the SV (arrowheads, Fig 6B; dark and bright red bars; Fig 6C). The distribution in the more distal zone (200-300 μm from SV) was similar to the control (Fig 6C). In addition, we observed a large number of single spermatid heads in the TE region. Together, these observations suggest that loss of Dlg1 in the SCCs disrupted NBs during cyst rotation inside the TE before sperm release.

**Fig 6.**
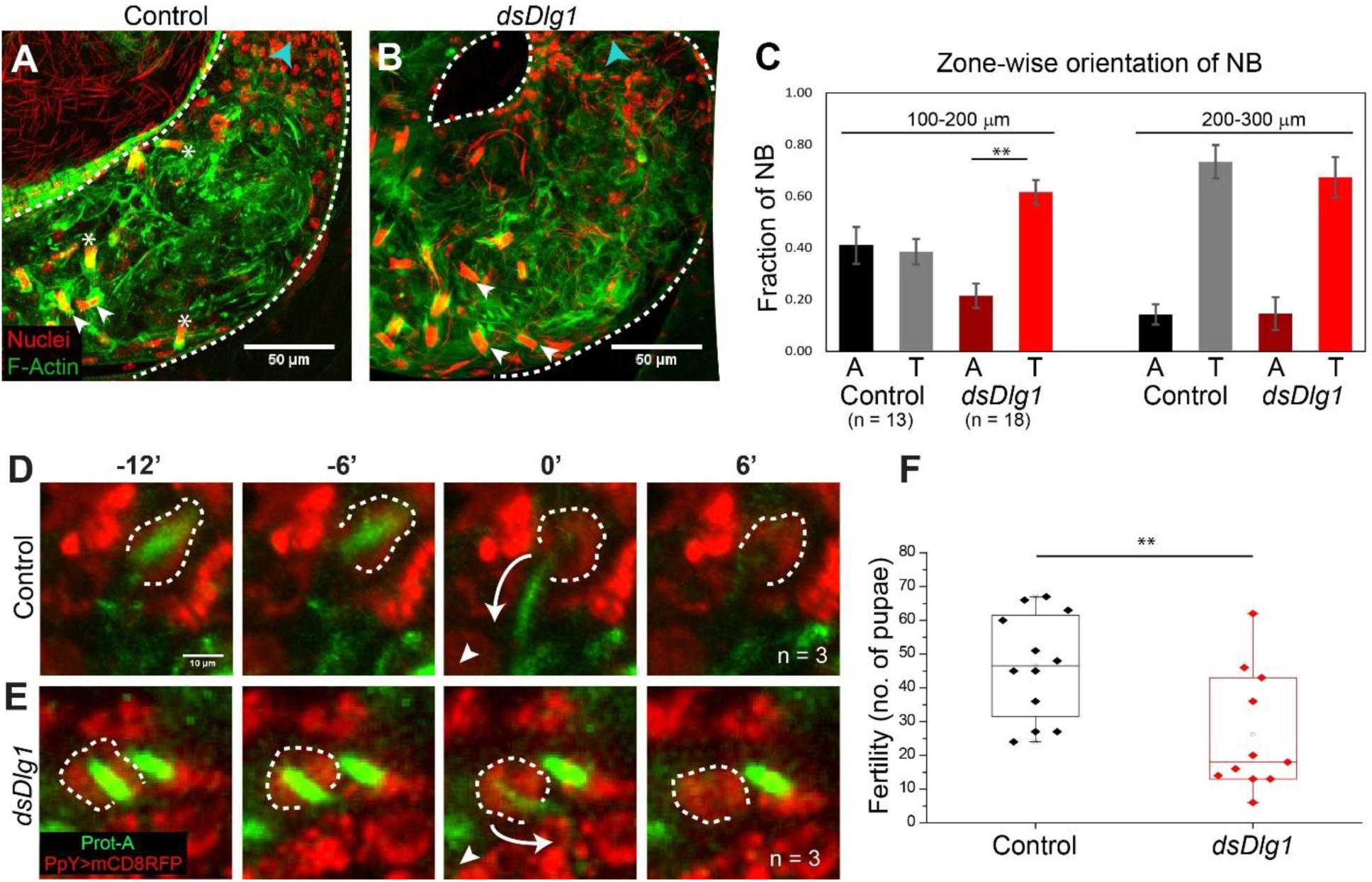
Dlg1 loss from the HCC-TCC interface at the coiling stage causes premature sperm release inside the testis. **A-B)** NB orientations in control **(A)** (*PpY*-*Gal4*> *dsGFP*) and *PpY*-*Gal4*> *dsDlg1* **(B)** testes. Testes were stained with the Hoechst dye (red) and Phalloidin (green), and the position of the actin cap was used as an indicator of whether the NBs were facing towards (white arrowhead) or away (asterisk) from the SV (direction of SV marked by blue arrowhead). Note the decrease in bundles facing away from SV in 100-200 μm region in **B.** **C)** Zone-wise distribution and the orientation of NBs (mean±s.e.m.) in the TE region. Distances from the SV were measured from the proximal end of the testicular duct. ‘A’ denotes the NB orientation away from SV, and ‘T’ denoted orientations towards the SV. P-value (^⋆⋆^<0.01) was calculated using the Mann-Whitney U test. Apart from these two classes a fraction of NBs was found with intermediate orientations that were not included in the graph. Spermatids exit testis in the ‘A’ orientation (Dubey et al., 2016). **D-E)** Time series from live imaging of the testes. Protamine A (green) marks spermatid head, while mCD8-RFP (red) marks cyst cell (dashed outline). In Control (*ProtA*-*GFP/UAS*-*Dicer; PpY*-*Gal4*> *UAS*-*mCD8 RFP*) **(D)**, spermatids heads are released (white arrow at time 0’) in the direction of the SV (indicated by white arrowhead). In the *dsDlg1* testes (*ProtA*-*GFP/UAS*-*dsDlg1*; *PpY*-*Gal4*>*UAS*-*mCD8 RFP*) **(E)**, spermatid heads are released in the TE region (white arrow at time 0’), even though the cyst has not turned to face away from the SV. **F)** Box plots depict the number of pupae produced by individual Control (*PpY*-*Gal4*> *dsGFP*) and *PpY*-*Gal4*> *dsDlg1* males in 24 hours. P-value (^⋆⋆^<0.01) was calculated using the Mann-Whitney U test.

To confirm the conjecture, we captured time-lapse images from whole testis *ex-vivo.* The spermatid heads were labeled with ProtamineA-GFP and cyst cell membrane was labeled with *PpY*-*Gal4*> *mCD8*-*RFP* in both the control (*UAS*-*Dicer*) and *dlg1* RNAi (*UAS*-*dsDlg1*) backgrounds. In the control testes, the NBs always retracted from the HCC during release in the direction of the SV (n = 3, Fig 6D; Movie S2). In the *dlg1* RNAi background, the spermatid heads retracted even though they were not facing away from the SV (n = 3, Fig 6E; movie S3). As a result, they were released prematurely inside the TE region. Hence, we conclude that the loss of Dlg1 in SCCs leads to loss of the SJs between the HCC and TCC, and that the integrity of this junction is critical to prevent cyst disruption and premature sperm release within the testis.

To understand the implications of the mechanical stability of the cyst enclosure during spermiation on male reproductive fitness, we carried out a fertility test. Individual males expressing the *GFP^dsRNA^* (control) and *Dlg1^dsRNA^* transgenes under *PpY*-*Gal4* were allowed to mate for 24 hours with three wild-type virgins each, and the number of pupae was counted. The results indicated that fertility of the *dlg1* RNAi males was significantly lower as compared to control males (Fig 6F). Together with the observation of the time-lapsed images, this result suggests that the premature release is detrimental to the male reproductive fitness.

## Discussion

### SJs between the somatic cyst cells are remodeled during spermatid maturation

The cyst capsule undergoes extensive changes in cell shape and size during the development of germ cells from the spermatogonial to coiled spermatid stages. The cyst has to maintain the enclosure to ensure proper differentiation of the germ cells. How the cyst manages to keep the enclosure intact during spermatogenesis was not known. Based on TEM observations, Tokuyasu reported the presence of septate junctions during late stages of *Drosophila* spermatogenesis (Tokuyasu et al., 1972). More recently, the SJ proteins Cora and Nrx-IV were shown to localize around the cyst during the early spermatogonial stages, and knockdown of these proteins at these stages led to an arrest in differentiation (Fairchild et al., 2015). In this study, we provided a systematic description of the septate junction morphogenesis during sperm development. Using Nrg-GFP as a marker, we showed that the SJs formed between the SCCs are dynamically rearranged towards the later stages before spermiation. During the spermatogonial stages, the SJ proteins are localized on both the germ cells and somatic cell membrane, while from the spermatocyte stages, these proteins are restricted to the SCCs. During the early elongated stages, the SJs proteins accumulate at the boundary between the HCC and TCC. This boundary localization remains until the coiled stages of spermatogenesis. Ultimately, the sperms release without breaching the SJ between the SCCs (Dubey et al., 2016), indicating that sperm release may take place due to a breach in the TCC.

A similar reorganization of SJ proteins has been described during *Drosophila* embryogenesis. It was shown that in epithelial cells of the trachea, until stage 13, SJ components localize all along the basolateral edges, and by stage 15 they are localized exclusively to the apico-lateral domain (Tiklová et al., 2010). Similarly, other studies have shown that until stage 15, Cora localizes all along the basolateral domain of the cells of the salivary gland, and post stage 15, they localize to the apico-lateral domain (Hall and Ward, 2016). Permeability experiments also suggest that the occluding property of SJs is attained by stage 15 (Paul et al., 2003). Ultrastructural studies show that mature SJs are formed by stage 16-17 (Tepass and Hartenstein, 1994). Together these observations suggested that a more diffuse localization of SJ components along the cell membrane is followed by the assembly of a compact, mature and functional occluding junction at a later stage.

In the testis, however, the SJ proteins are localized on both the germline and somatic cell membrane during the spermatogonial stages, and the somatic permeability barrier is established from the 4-cell stage. Loss of the SJ components-Dlg1, Nrx-IV, and Cora-from the germ-soma interface during this period disrupted the permeability barrier and affected subsequent differentiation to the spermatocyte stage (Fairchild et al., 2015; Gupta et al., 2018). A similar loss of permeability due to the knockdown of Armadillo/β-catenin, however, did not appear to affect the immediate differentiation to the spermatocyte stage (Gupta et al., 2018). Therefore, the SJ proteins Dlg1, Cora, and Nrx-IV are likely to regulate the germline differentiation independent of their role in establishing the barrier function. Due to the distinctive morphogenetic profiles of the cellular interfaces, one could also recognize that the SJ proteins relocalize after meiosis to a new interface between the SCCs during spermatid elongation and this association is retained all through the remaining period of spermatid differentiation.

### Atypical SJs forms only after meiosis and during spermatid elongation

In concurrence with a previous report (Tokuyasu et al., 1972), the TEM data also suggested that a ladder-like, septate pattern is formed after the elongation stages, and we do not find the presence of SJs during mitotic and meiotic stages. In the mammalian testis, the BTB is formed after the mitotic stages, and it serves to provide an isolated environment to the meiotic and post meiotic population (Mruk and Cheng, 2015). We could not find any SJ-like feature between the germ cells or at the germ-soma interface during the early stages. The first electron-dense material appeared at the interface of the SCCs encapsulating elongated spermatids. Therefore, SJs are unlikely to contribute to establishing the permeability barrier during the spermatogonial stages.

After the reorganization, SJ proteins localize at the HCC-TCC interface, which is further compacted during spermatid individualization. Classically, TJs and SJs localize in a tight band at the apicolateral domains of an epithelium, thereby stitching the neighboring cells together. In comparison, the SJs formed between two cyst cells (HCC and TCC) is extended along the entire cellular interface. In this way, they seal the enclosure formed by the head-to-head association of two SCCs, which is distinct from the interactions established by these junctions in a monolayer. Due to this unusual arrangement, we call this an atypical septate junction. The junction moves from the middle of the elongated cyst towards the base of the NB during the coiling stages. This kind of the extensive morphogenesis is unique to *Drosophila* testis. It indicated substantial morphological restructuring of the HCC and TCC during this period. Previously, TEM analysis of testis sections suggested that the movement of the junction and reshaping of the cyst cells may occur during individualization or early coiling stage (Tokuyasu et al., 1972). Our results obtained from time-lapse imaging of live testis preparations support this hypothesis and provides experimental proof for the repositioning of the junction.

### SJs between the Head and Tail cyst cells provide mechanical stability to the somatic enclosure during spermatid coiling

Although *PpY*-*Gal4* is expressed in the SCCs from the post-mitotic stages (Jung et al., 2007), the *PpY*-*Gal4*-mediated expression of *dsDlg1* could only eliminate the protein from the SCCs at the last stage of spermatid maturation, when the cyst entered the TE. It increased the propensity of premature spermatid release inside the testis. Time-lapse analysis indicated that these releases occurred at an unusual orientation. Together, these observations suggest that the turning of the cyst is a mechanically stressful event which can only be accomplished if the cyst cells are tightly adherent. SJs are classically thought to provide a fluid access barrier across an epithelium. However, the evidence from *Drosophila* testis could also suggest a role in providing mechanical stability. In mice testis, knockdown of claudin-11 led to sloughing off the Sertoli cells from the seminiferous tubule, indicating that the TJs are required to maintain structural integrity in an epithelium (Mazaud-Guittot et al., 2010). Recently, it has been seen in *Xenopus* embryos that loss of TJ proteins leads to an increase in tension on Adherens junctions during cytokinesis (Hatte et al., 2018). Therefore, apart from serving the barrier properties, SJs and TJs may also help in generating tissue resistance to mechanical strain which is essential for maintain organ shape and integrity during development and in adult stages.

## Materials and Methods

### *Drosophila* stocks and culture conditions-

All *Drosophila* stocks and crosses were maintained at standard cornmeal *Drosophila* medium at 25 °C. Freshly emerged flies are separated from females and were allowed to age for 2-4 days before dissection. For RNAi experiments, the freshly emerged males were kept at 28 °C to increase the penetrance of the RNAi. The list of the stocks and their sources are listed in the supplemental Table S1. We thank the *Drosophila* community for their generous gift of the fly stocks.

### Fertility assay

Each freshly emerged, *PpY*-*Gal4*> *dsGFP* (control) and *PpY*-*Gal4*> *dsDlg* males were kept with three Canton-S females for four days at 28 °C to allow for accumulated sperm to be cleared out. On the fourth day, each of these males was extracted and mated with three fresh virgin females (Canton-S) for 24 hours at 28 °C in separate vials. Then all the flies were discarded. Subsequently, the number of pupae in the vial were counted as a measure of the male fertility.

### Immunostaining

For whole mount immunostaining, the testes were dissected in 1X PBS followed by fixation in 4% Para-formaldehyde (PFA) for 30 mins-1 hour at room temperature. Post-fixation, testes were washed with PTX (0.3% Triton-X in PBS), three times, 10 mins each. After washing, the samples were incubated with Primary antibody diluted in PTX overnight at 4 °C. The primary antibody solution was washed off with PTX, and samples were incubated with Alexa-dye tagged secondary antibodies (Invitrogen) for 2-4 hours at room temperature. After washing, samples were stained with 0.001% Hoechst 33342 (Sigma Chemical Co. USA), and 10 μM Phalloidin-Atto568/647 (Sigma Chemical Co. USA) for 30 mins, washed and mounted in Vectashield^®^ mounting medium (Vector Laboratory Inc., USA) on a glass slide. For testis squash preparation, the testes were dissected and kept in 50 μl of PBS on a glass slide. A coverslip was placed on top of the sample and gently pressed against the slide. Extra PBS was removed, and the slide was plunged into liquid nitrogen for two minutes. After removing the coverslip, making sure the sample remains on the slide, the slide was then incubated with 95% ethanol, followed by fixation in 4% PFA for an hour. Further processing is same as described for whole mount immunostaining. The primary antibodies used are as follows: Anti-Dlg1 (4F3, DSHB; 1:100), Anti-Cora (C615.16, DSHB; 1:100), Anti-Vasa (DSHB; 1:50), Alpha-spectrin (3A9, DSHB; 1:100), Anti-Eya (eya10H6, DSHB; 1:100) and Anti-tj (Dorothea Godt, University of Toronto, Canada; 1:1000).

### Transmission electron microscopy

Three to Four days old CantonS flies were dissected in 1X PBS and fixed for 4-6 hours, with Karnovsky’s fixative (pH-7.4) at room temperature. Samples were washed with 100 mM phosphate buffer (pH-7.4), then post-fixed with K_2_Cr_2_O_7_–OsO_4_ mixture for 2 hours on ice, which was followed by 1-hour incubation at room temperature. After 3 to 5 washes in 100 mM phosphate buffer, the specimens were dehydrated in a graded series of ethanol and propylene oxide. Finally, they were embedded in Durcupan (Fluka, Electron Microscopy Sciences, USA) epoxy resin mix prepared according to the manufacturer protocol and polymerized at 60°C overnight. Specimens were then sectioned with a glass and diamond knife on LEICA-EM-UC6 (Leica Microsystems, Germany). Ultrathin sections were collected on Formvar-carbon coated copper slots. These sections were examined on Libra120EFTEM Transmission Electron microscope (Carl Zeiss AG, Germany).

### Image Analysis and Quantification

Images were acquired using Olympus FV1000SPD and FV3000SPD Laser scanning confocal microscopes (Olympus Co., Japan). Live Imaging was performed on FV1000SPD, as described by (Dubey et al., 2016). The images were analyzed using Fiji-ImageJ (http://fiji.sc/Fiji). The pair-wise significance of difference (p-value) was estimated using the Mann-Whitney U-test.

## Acknowledgments

We thank Lalit Borde for the help with the TEM imaging; Prof. John Belote, Syracuse University, NY, USA; Prof. Benny Shilo, Weizmann Inst., Israel, and Dr. Dorothea Godt, University of Toronto, Canada for reagents. We acknowledge the Fly facility at the National Centre for Biological Sciences (NCBS), Bangalore, India; Bloomington Drosophila Stock Centre (BDSC), Indiana USA; and Vienna Drosophila Resource Centre (VDRC), Austria; for fly stocks; and Developmental Studies Hybridoma Bank, Iowa for antibodies. We also thank Prof. M. Narasimha and KR lab members for the assistance with various reagents and stocks. The research was supported by an intramural grant of TIFR, DAE, Govt. of India.

## Author contributions

PD and KR conceived the project and planned experiments. PD and TK carried out the experiments and compiled the figures. SG and SS contributed the TEM data. KR wrote the manuscript with help from PD and TK.

## References

Banerjee, S., Sousa, A. D. and Bhat, M. A. (2006). Organization and Function of Septate Junctions. 46, 65–77.

Baumgartner, S., Littleton, J. T., Broadie, K., Bhat, M. A., Harbecke, R., Lengyel, J. A., Chiquet-ehrismann, R., Prokop, A., Bellen, H. J. and Miescher-institute, F. (1996). A Drosophila Neurexin Is Required for Septate Junction and Blood-Nerve Barrier Formation and Function. 87, 1059–1068.

Cheng, C. Y. and Mruk, D. D. (2012). The Blood-Testis Barrier and Its Implications for Male Contraception. Pharmacol. Rev. 64, 16–64.

Dubey, P., Shirolikar, S. and Ray, K. (2016). Localized, Reactive F-Actin Dynamics Prevents Abnormal Somatic Cell Penetration by Mature Spermatids. Dev. Cell 38, 507–521.

Fabrizio, J. J., Boyle, M. and DiNardo, S. (2003). A somatic role for eyes absent (eya) and sine oculis (so) in Drosophila spermatocyte development. Dev. Biol. 258, 117–128.

Fairchild, M. J., Smendziuk, C. M. and Tanentzapf, G. (2015). A somatic permeability barrier around the germline is essential for Drosophila spermatogenesis. Development 142, 268–281.

Fehon, R. G., Dawson, I. A. and Artavanis-Tsakonas, S. (1994). A Drosophila homologue of membrane-skeleton protein 4.1 is associated with septate junctions and is encoded by the coracle gene. Development 120, 545–557.

Furuse, M., Fujita, K., Hiiragi, T., Fujimoto, K. and Tsukita, S. (1998a). Claudin-1 and -2: novel integral membrane proteins localizing at tight junctions with no sequence similarity to occludin. J. Cell Biol. 141, 1539–50.

Furuse, M., Sasaki, H., Fujimoto, K. and Tsukita, S. (1998b). A single gene product, claudin-1 or -2, reconstitutes tight junction strands and recruits occludin in fibroblasts. J. Cell Biol. 143, 391–401.

Ghosh-Roy, A., Kulkarni, M., Kumar, V., Shirolikar, S. and Ray, K. (2004). Cytoplasmic Dynein – Dynactin Complex Is Required for Spermatid Growth but Not Axoneme Assembly in. 15, 2470–2483.

Ghosh-Roy, A., Desai, B. S., Ray, K. (2005). Dynein Light Chain 1 Regulates Dynamin-mediated F-Actin Assembly during Sperm Individualization in Drosophila. 16, 3107–3116.

Griswold, M. D. (1998). The central role of Sertoli cells in spermatogenesis. 9, 411–416

Gupta, S., Varshney, B., Chatterjee, C., Ray, K. (2018) Somatic ERK activation during the transit amplification is essential for maintaining the synchrony of germline divisions in *Drosophila* testis. O. Bio (in press)

Hall, S. and Ward, R. E. (2016). Septate Junction Proteins Play Essential Roles in Morphogenesis Throughout Embryonic Development in Drosophila. G3-Genes Genomes Genet. 6, 2375–2384.

Hartsock, A. and Nelson, W. J. (2008). Adherens and Tight Junctions: Structure, Function and Connection to the Actin Cytoskeleton. Biochim Biophys Acta 1778, 660–669.

Hatte, G., Prigent, C. and Tassan, J.-P. (2018). Tight junctions negatively regulate mechanical forces applied to adherens junctions in vertebrate epithelial tissue. J. Cell Sci. 131, jcs208736.

Hudson, A. G., Parrott, B. B., Qian, Y. and Schulz, C. (2013). A Temporal Signature of Epidermal Growth Factor Signaling Regulates the Differentiation of Germline Cells in Testes of Drosophila melanogaster. PLoS One 8: e70678

Jung, A., Hollmann, M. and Schafer, M. A. (2007). The fatty acid elongase NOA is necessary for viability and has a somatic role in Drosophila sperm development. J. Cell Sci. 120, 2924–2934.

Leatherman, J. L. and Di Nardo, S. (2008). Zfh-1 controls somatic stem cell self-renewal in the Drosophila testis and nonautonomously influences germline stem cell self-renewal. Cell Stem Cell 3, 44–54.

Li, M. A., Alls, J. D., Avancini, R. M., Koo, K. and Godt, D. (2003). The large Maf factor traffic jam controls gonad morphogenesis in Drosophila. Nat. Cell Biol. 5, 994–1000.

Lindsley, D. I. and Tokuyasu, K. T. (1980). Spermatogenesis. In Genetics and Biology of Drosophila, 2nd edn (ed. M. Ashburner and T. R. Wright), pp. 225–294. New York: Academic Press.

Locke, M. (1965). The structure of septate desmosomes. J. Cell Biol. 25, 166–169.

Mazaud-Guittot, S., Meugnier, E., Pesenti, S., Wu, X., Vidal, H., Gow, a and Le Magueresse-Battistoni, B. (2010). Claudin 11 deficiency in mice results in loss of the Sertoli cell epithelial phenotype in the testis. Biol. Reprod. 82, 202–213.

Mruk, D. D. and Cheng, C. Y. (2015). The mammalian blood-testis barrier: Its biology and regulation. Endocr. Rev. 36, 564–591.

Nelson, K. S., Furuse, M. and Beitel, G. J. (2010). The Drosophila claudin Kune-kune is required for septate junction organization and tracheal tube size control. Genetics 185, 831–839.

Papagiannouli, F. (2013). The internal structure of embryonic gonads and testis development in Drosophila melanogaster requires scrib, lgl and dlg activity in the soma. Int. J. Dev. Biol. 57, 25–34.

Papagiannouli, F. and Mechler, B. M. (2009). discs large regulates somatic cyst cell survival and expansion in Drosophila testis. Cell Res. 19, 1139–49.

Paul, S. M., Ternet, M., Salvaterra, P. M. and Beitel, G. J. (2003). The Na + / K + ATPase is required for septate junction function and epithelial tube-size control in the Drosophila tracheal system. Development. 130, 4963–4974.

Tepass, U. and Hartenstein, V. (1994). The development of cellular junctions in the Drosophila embryo. Dev. Biol. 161, 563–596.

Tiklová, K., Senti, K. A., Wang, S., GräCurrency Signslund, A. and Samakovlis, C. (2010). Epithelial septate junction assembly relies on melanotransferrin iron binding and endocytosis in Drosophila. Nat. Cell Biol. 12, 1071–1077.

Tokuyasu, K. T., Peacock, W. J. and Hardy, R. W. (1972). Dynamics of spermiogenesis in Drosophila melanogaster. II. Coiling process. Zeitschrift für Zellforsch. und Mikroskopische Anat. 127, 492–525.

White-cooper, H. (2004) Spermatogenesis: analysis of meiosis and morphogenesis. In: Henderson D, editor. Methods in molecular biology. Totowa, NJ: Humana Press. pp45–75. doi: 10.1385/1-59259665-7:45-

Woods, D. F. and Bryant, P. J. (1989). Molecular cloning of the lethal(1)discs large-1 oncogene of Drosophila. Dev. Biol. 134, 222–235.

Woods, D. F., Hough, C., Peel, D., Callaini, G. and Bryant, P. J. (1996). Dig Protein Is Required for Junction Structure, Cell Polarity, and Proliferation Control in. 134, 1469–1482.

Wu, V. M., Schulte, J., Hirschi, A., Tepass, U. and Beitel, G. J. (2004). Sinuous is a Drosophila claudin required for septate junction organization and epithelial tube size control. J. Cell Biol. 164, 313–323.

J. Xu, F. Anuar, S. M. Ali, Y. N. Mei, D. C. Y. Phua, and W. Hunziker. (2009). Zona occludens-2 is critical for blood-testis barrier integrity and male fertility. Molecular Biology of the Cell, vol. 20, no. 20, pp. 4268–4277, 2009.

Zoller, R. and Schulz, C. (2012). The Drosophila cyst stem cell lineage: Partners behind the scenes? Spermatogenesis 2, 145–157.

